# Deep topographic proteomics of a human brain tumour

**DOI:** 10.1101/2022.03.21.485119

**Authors:** Simon Davis, Connor Scott, Janina Oetjen, Philip D Charles, Benedikt M Kessler, Olaf Ansorge, Roman Fischer

## Abstract

The spatial organisation of cellular protein expression profiles within tissue determines cellular function and is key to understanding disease pathology. To define molecular phenotypes in the spatial context of tissue, there is a need for unbiased, quantitative technology capable of mapping proteomes within tissue structures. Here, we present a workflow for spatially-resolved, quantitative proteomics of tissue that generates maps of protein abundance across tissue slices derived from a human atypical teratoid-rhabdoid tumour (AT/RT). We employ spatially-aware algorithms that do not require prior knowledge of the fine tissue structure to detect proteins and pathways with spatial abundance patterns. We identified PYGL, ASPH and CD45 as spatial markers for tumour boundary and reveal immune response-driven, spatially-organised protein networks of the extracellular tumour matrix. Overall, this work informs on methods for spatially resolved deep proteo-phenotyping of tissue heterogeneity, to push the boundaries of understanding tissue biology and pathology at the molecular level.

## Main

Tissues contain various microscopic features, cell types, and phenotypically diverse subpopulations while the location of cells within a tissue and their spatial neighbourhood is crucial for determining their identity and function^1–6^. The cellular composition of tissue has substantial influence on measured co-expression signals within the molecular profiles of bulk tissue and their microenvironment, contributing to the influence of cellular function, signalling and different disease outcomes^7–11^. Recent technology developments in spatially-resolved sequencing technologies have enabled the characterisation of spatially heterogeneous gene expression profiles within tissues^12,13^. However, while genomic and transcriptomic alterations act as drivers of disease, the proteins they encode regulate essentially all cellular processes and therefore also need consideration when investigating tissue spatial heterogeneity^14^.

A range of mass spectrometry (MS)-based techniques are available to map the distribution of proteins throughout tissues and cells. Mass spectrometry imaging (MSI) enables the determination of proteins or other molecules within a sample by rastering an ion source over a sample in a grid pattern. MSI techniques to visualise bio-molecules *in situ* cannot generate in-depth proteome data and may require prior knowledge of measurement targets^15–18^.

Laser capture microdissection (LCM) is well-placed to address the limitations of the spatially-resolved mass spectrometry methods described above^19^. LCM can extract regions from a tissue slice ranging from single cells to square millimetres of tissue^20,21^. We and others have previously described several methods coupling LCM to MS-based proteomics^22– 26^. The approach has been used to investigate a wide range of tissue biology, generally in a ‘feature-driven’ approach, extracting tissue regions based on visible features^27–30^. Mund et al. recently developed the concept of Deep Visual Proteomics by combining high-resolution imaging and machine learning-based image analysis to classify, isolate and analyse cells with a sensitive proteomics workflow^31^.

However, sampling in a systematic manner, like MSI, could reveal novel tissue fine structure and give insights into the spatial protein expression patterns. For example, Piehowski et al. used LCM-proteomics to sample mouse uterine tissue in a rastered grid with a resolution of 100 µm and their custom, robotic, nanolitre-scale nanoPOTS sample preparation platform to quantify over 2,000 proteins within the tissue^32^.

Here, we systematically performed spatially-resolved measurements of a human brain tumour proteome using laser capture microdissection to a depth of over 5,000 proteins. We use spatially-aware statistical methods to identify proteins and pathways displaying differential spatial and clustered expression within tissue sections. This did not require prior knowledge of tissue structures, features, or pathology. Furthermore, clustering of protein expression reveals new, spatially defined proteo-phenotypes within the otherwise homogeneous macrostructure of the analysed tumour revealing spatially resolved ECM biology correlating with immune response.

## Results

In order to establish required tissue areas to achieve a target proteome depth of 4000 quantified proteins in low (Orbitrap Fusion Lumos, 60-gradient) and medium throughput proteomics platforms (TimsTOF Pro, 17-minute gradient) we collected tissue areas ranging from 316 µm^2^ to 1,000,000 µm^2^ from 10 µm-thick sections of human brain (Figure S1). We observed that areas above 316,000 µm^2^ result in diminishing returns with protein identifications scaling between 282 and 3480 on the Orbitrap and 127 and 3318 on the timsTOF Pro platforms, respectively.

### Proteomic topography of a human brain tumour

After characterising the upper and lower limits of the workflow, we sampled a 10 µm thick section of an atypical teratoid-rhabdoid tumour (AT/RT) block (∼20 × 15 mm). The tissue was subdivided into 384 (24 × 16) square ‘pixels’ with an area of ∼694,000 µm^2^ (side length of 833 µm); each pixel was isolated by LCM and processed with our LCM-SP3 protocol^24^ (Figure 1). Each sample was analysed on the 17-minute timsTOF Pro setup. In total, 5,321 proteins were identified, with 32 – 4,741 proteins identified per sample (Figure S2). This range includes empty pixels where no tissue was visible, demonstrating a low level of contamination throughout the workflow. Quantified proteins can be mapped back to their original positions within the tissue grid. Figure 2A shows proteomic maps for four example proteins, liver glycogen phosphorylase (PYGL), peripherin (PRPH), haemoglobin (HBB) and histone H4 (HIST1H4A). Glycogen phosphorylase releases glucose from glycogen for entry into glycolysis, and its expression in cancer is associated with malignant phenotypes, hypoxia resistance and cancer cell survival^33^. Peripherin is an intermediate filament protein without a clear function and is highly expressed during development and after nerve injury; its expression pattern is consistent with the tumour growth into surrounding normal brain tissue^34–36^.

**Figure 1.**
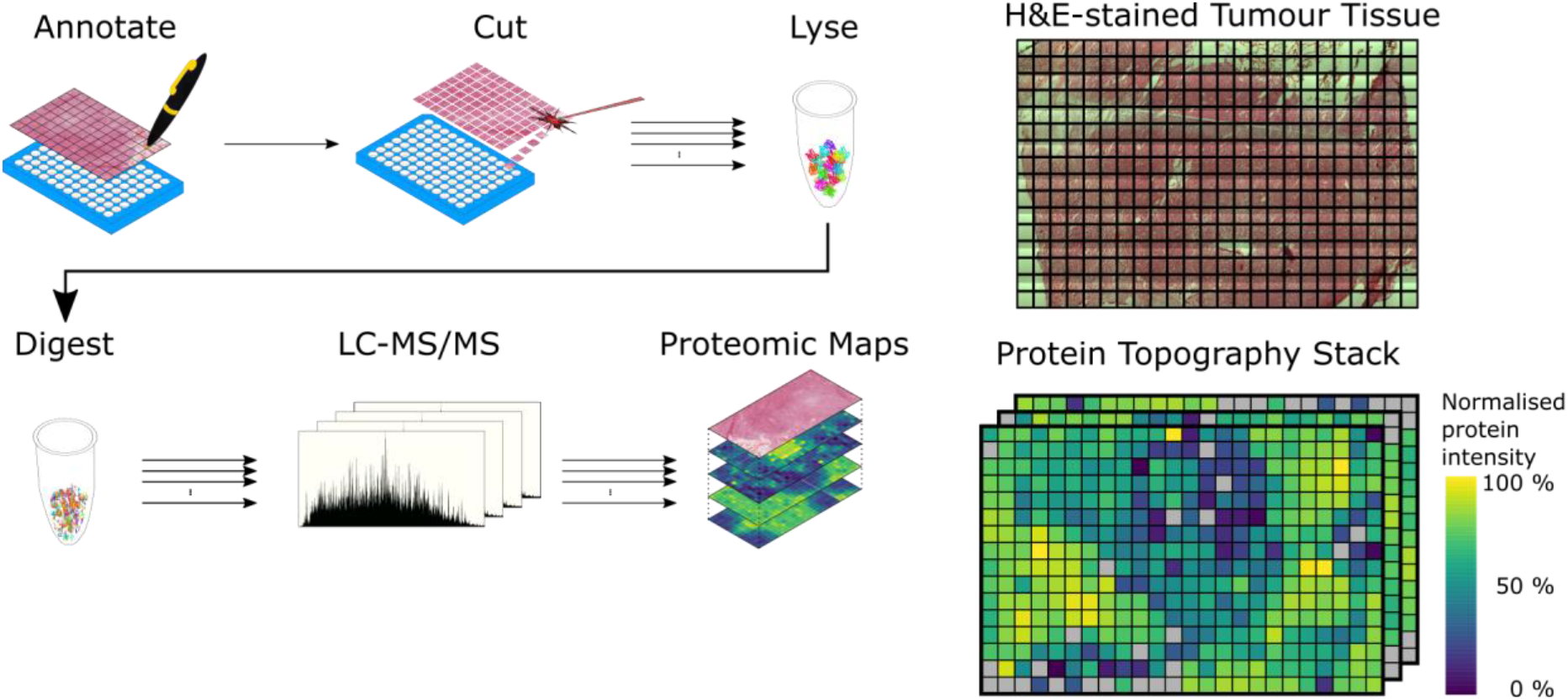
Overview of the spatially resolved proteomics workflow. Tissue is segmented into a regular grid shape, and each element of the grid is isolated by laser capture microdissection (LCM). Proteins from each individually lysed sample are digested before analysis by LC-MS/MS. The quantitative information for each protein can be mapped back to its location within the gridded tissue and visualised in a topographic protein map, with one map per protein quantified.

**Figure 2.**
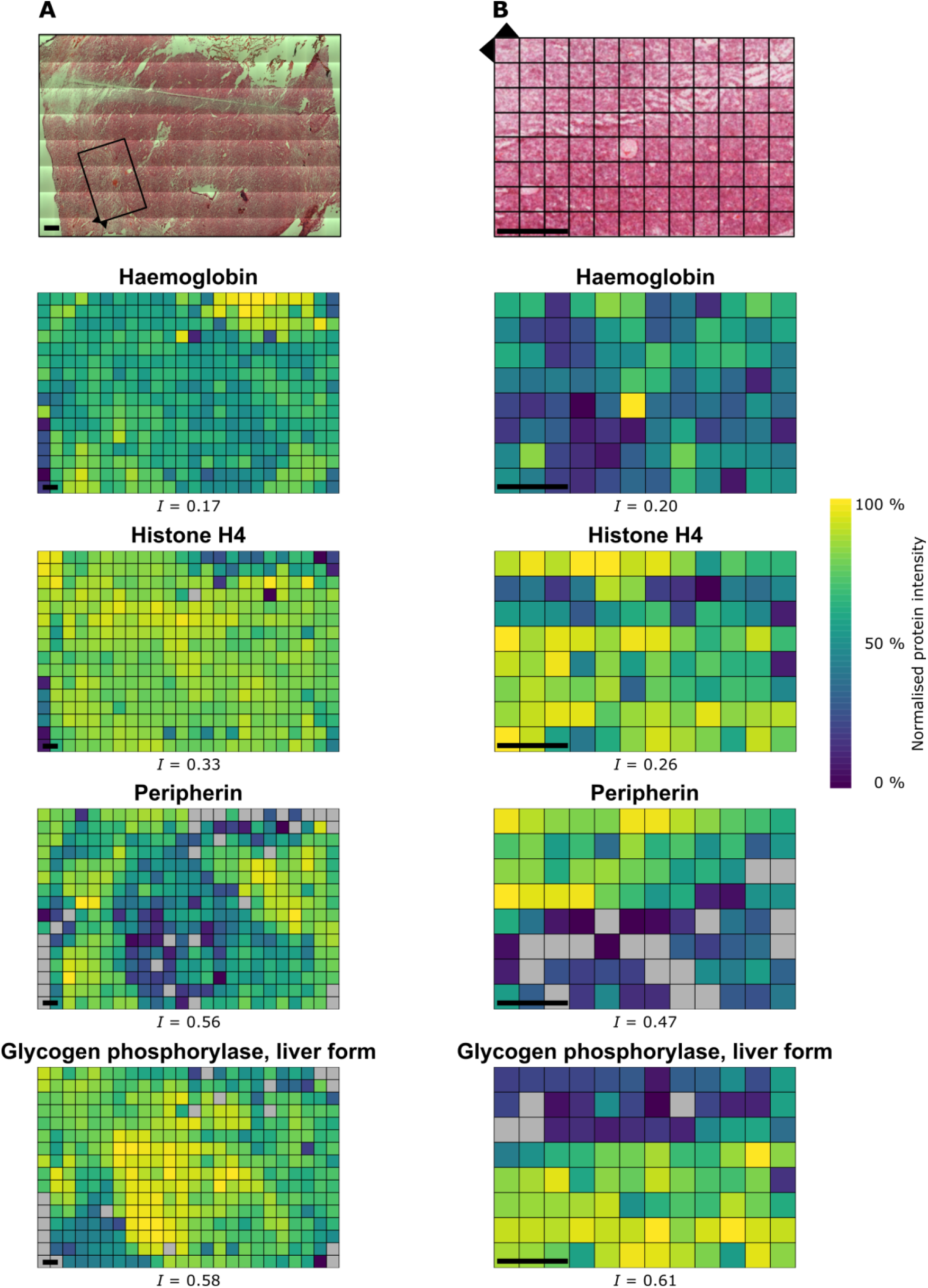
Spatial proteomic maps of AT/RT tumour tissue reveal regional boundaries. Normalised protein intensity of four example proteins mapped back to the original spatial positions within the atypical teratoid-rhabdoid tumour (AT/RT) tumour tissue at a resolution of 833 µm (A) and 350 µm (B) with their corresponding Moran’s Index of spatial autocorrelation (*I*). Box in (A) represents the area analysed in an adjacent tissue section (B). Scale bar = 1 mm. Normalised protein intensities are scaled separately for each protein. Grey = not detected.

We tested the 4,306 proteins quantified in at least 9 pixels for spatial autocorrelation^37^ using the unsupervised Moran’s *I* test. Moran’s I returns values between -1 for complete dispersion and +1 for complete correlation, with 0 indicating random distribution. Removing empty pixels and areas of haemorrhage from the analysis, we found that 3,212 proteins demonstrated correlated spatial expression profiles with (*q* ≤ 0.05), while excluding any impact of sampling and processing on spatial variability (Figure S3).

In order to increase spatial resolution particularly in the ‘brain/tumour’ interface as identified by H&E staining, we sampled with 96 pixels with 350 µm x 350 µm side-length. This region of tissue contains predominantly normal and neoplastic cells along with a large, prominent blood vessel. In total, 3,994 proteins were quantified in at least one pixel. The increased resolution proteomic maps for PYGL, PRPH, haemoglobin and histone H4 are shown in Figure 2B. Their expression is consistent with the large field-of-view data, with PYGL and PRPH showing opposite expression patterns across the margin between solid tumour (high PYGL, low PRPH) and brain/tumour interface (low PYGL, high PRPH) and haemoglobin co-localising with the visible blood vessels. Histone H4 shows even expression across the two annotated areas, with a region of lower expression corresponding with a visibly diffuse patch of tissue. Of the 3,050 proteins quantified in at least 9 pixels, 1,375 show evidence for significant spatial autocorrelation (Moran’s *I* test, *q* ≤ 0.05).

Three candidate proteins showing significant spatial variation were selected for follow-up immunohistochemistry (IHC) staining: glycogen phosphorylase, aspartate beta-hydroxylase (ASPH) and CD45 (PTPRC) to validate the spatially resolved protein expression data generated above. The IHC staining images closely resemble the protein intensity distributions (Figure 3A, C, E) within the proteomic maps for these three proteins. Both PYGL and ASPH show intense IHC staining in the region of solid tumour (Figure 3B,D), and CD45 shows intense staining in the region of tissue corresponding to the upper-left pixels in the proteomic map (Figure 3F).

**Figure 3.**
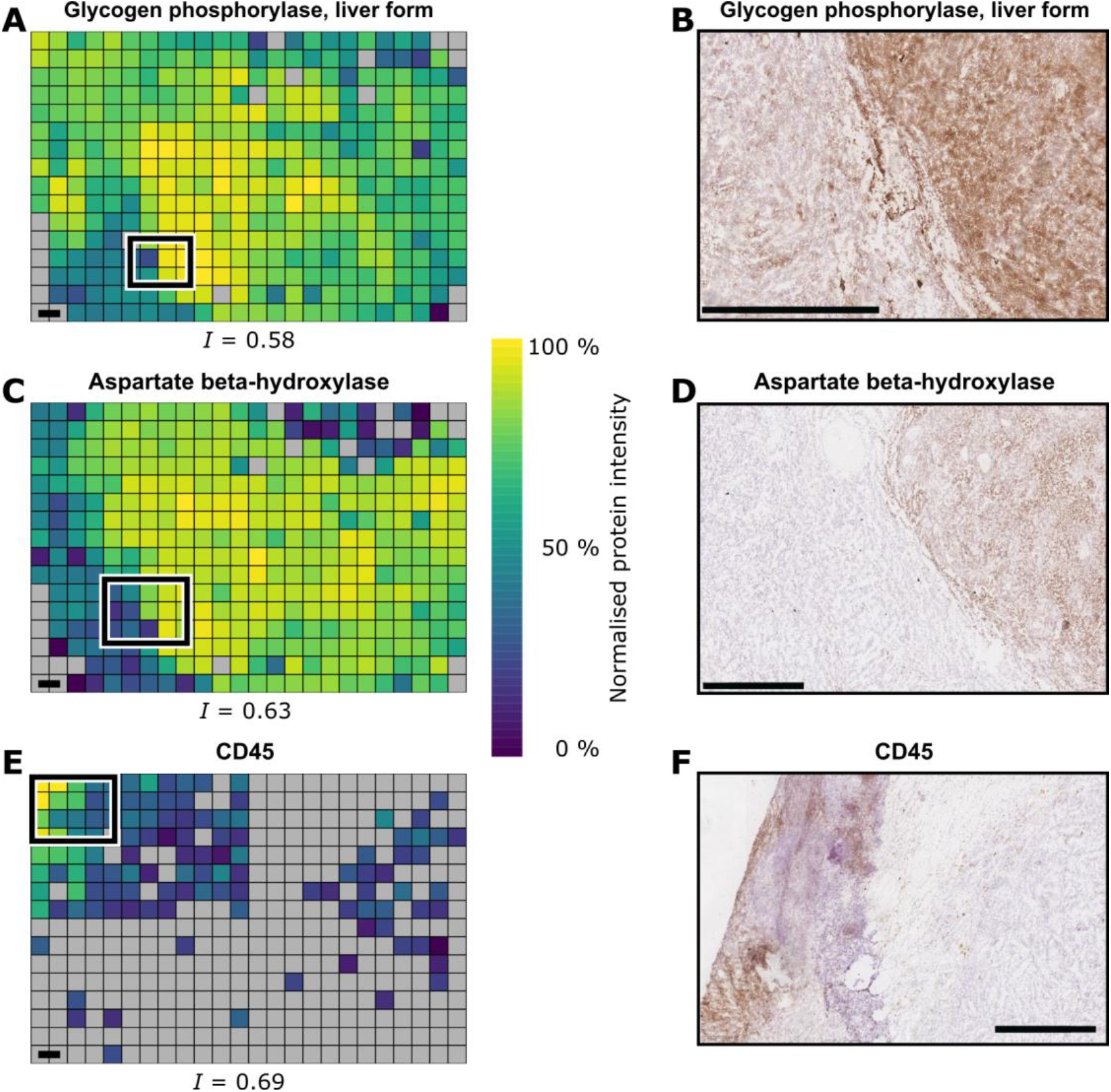
Immunohistochemistry validation of AT/RT proteomic maps. Proteomic maps of proteins targeted for follow-up IHC staining and IHC images. (A,C,E) Normalised protein intensity maps with their corresponding Moran’s Index of spatial autocorrelation (*I*). Normalised protein intensities are scaled separately for each protein. Grey = not detected. Rectangles depict the approximate location displayed in IHC images. (B,D,F) AT/RT tissue stained and visualised by IHC. All scale bars = 1 mm.

### Spatial proteomic mapping highlights molecular pathways underlying tissue heterogeneity

As Moran’s *I* measures global autocorrelation, it does not indicate where the locations that drive the autocorrelation occur. To investigate which regions of the sampled tissue show similar expression, the data were clustered and the cluster labelling of pixels were mapped back to their spatial location (Figure 4A). The clusters generally form contiguous regions in space, with some long-range co-clustering in smaller clusters. The margin between solid tumour and brain/tumour interface is well-represented by the border between cluster 1 (solid tumour) and cluster 3 (brain/tumour interface). In addition to the cluster map, the assigned clusters were plotted onto a uniform manifold approximation and projection (UMAP) visualisation (Figure 4B). The clusters visible in the UMAP plot correspond well to the spatial clusters in Figure 4A. This clustering approach generates spatially well-defined clusters, allowing for a feature-driven approach without prior knowledge of the histopathological details as the clustering is performed only on the protein quantitation data, without information on the spatial relationship between samples. Proteins quantified were tested for significantly differing intensities across the clusters using a one-way ANOVA test. There are 3,512 proteins with significant evidence (q ≤ 0.05) for differential intensity between two or more clusters.

**Figure 4.**
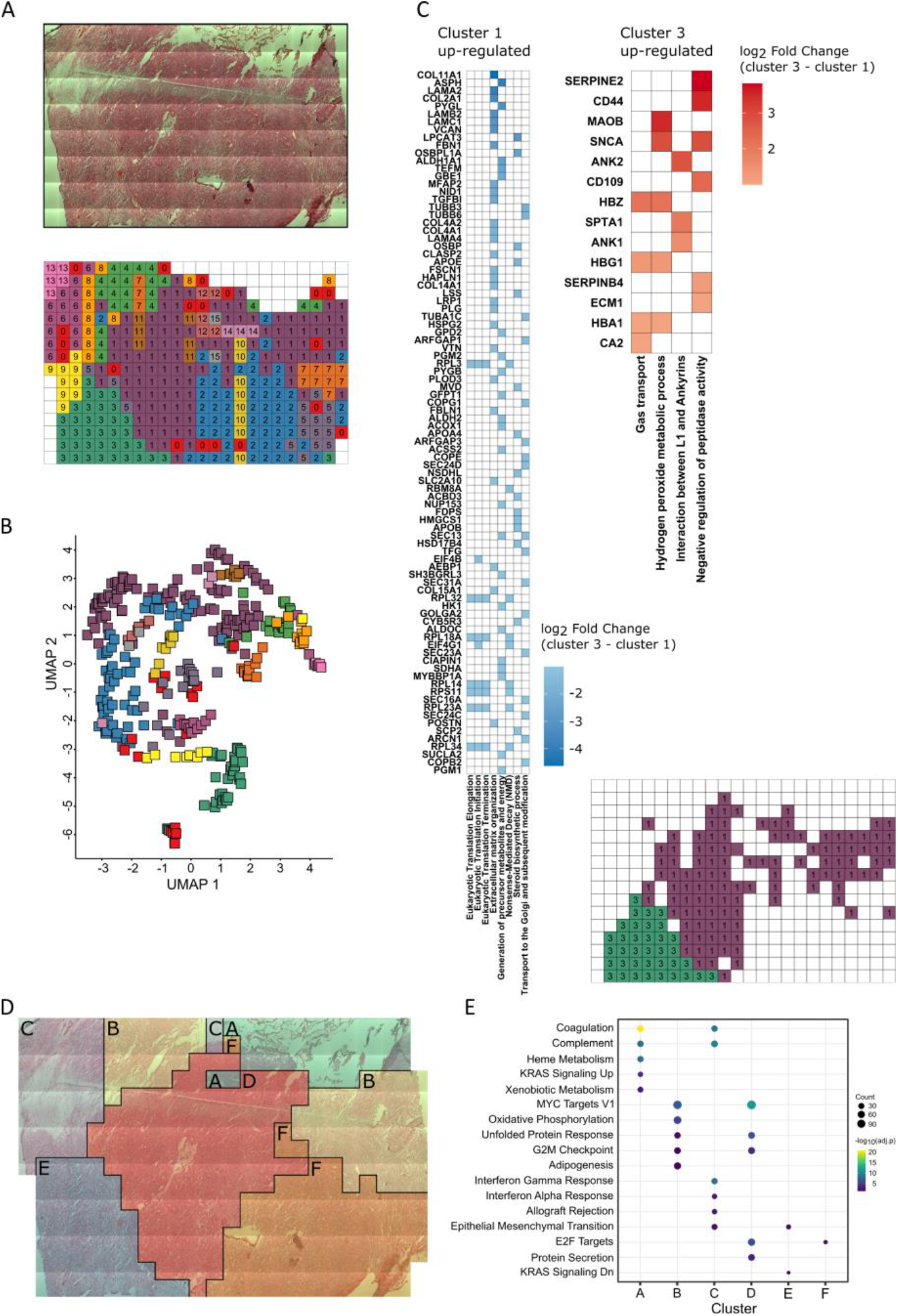
Clustering and functional analysis displays regional AT/RT proteomic maps in the tumour microenvironment. (A) Map of cluster assignment based on hierarchical clustering and the dynamic tree cut algorithm. This clustering is spatially-unaware and is based only on protein abundance values. White pixels represent excluded empty regions and regions of haemorrhage (B) UMAP embedding of data coloured by cluster assignment in (A). (C) GSEA of clusters 1 & 3 from (A). Gene set membership is indicated by colouring the cell with that protein’s log_2_ fold-change. (D) Cluster-map of ATRT tissue generated by affinity network fusion (ANF). This clustering is spatially-aware, meaning the relative spatial locations of samples and protein abundance values are used to generate these clusters. (E) Enriched MSigDB Hallmark gene sets within marker proteins of the clusters shown in the ANF cluster-map. Significantly enriched hallmarks for each cluster are indicated by the presence of circles. The size and colour of the circles represent the number of proteins contributing to that term and the adjusted *p* value of the enrichment, respectively.

Pairwise Tukey *post-hoc* tests were used to generate fold-change estimates between pairs of clusters, and these fold-changes were used for gene set enrichment analysis (GSEA). Functional analysis of clusters 1 & 3 indicates that proteins and processes involved in protein translation and modification, extracellular matrix organisation, energy pathways, mRNA processing, steroid synthesis, neuronal cell adhesion, and neuronal differentiation are differentially abundant between the region of solid tumour and brain/tumour interface (Figure 4C).

In order to discover new spatial relationships between phenotypically similar and distinct areas, we performed unbiased spatial clustering by integrating a proteomic similarity matrix (rank correlation of top 25 % most variable proteins) with a complementary spatial similarity network (based on Euclidean distances) using Affinity Network Fusion (ANF)^38^. This spatially-aware clustering method resulted in six clusters (Cluster A-F) covering the regions of solid tumour, brain/tumour interface, immune infiltration, and haemorrhage (Figure 4D).

The functional differences between these clusters were investigated by first determining ‘marker proteins’ for each cluster by testing for differential abundance of a protein in one cluster versus all other clusters, iteratively for each protein-cluster pair. Protein markers for each cluster were then tested for functional enrichment against the MSigDB Hallmark gene sets (Figure 4E)^39^. Cluster A shows enrichment for blood-related hallmarks, consistent with this cluster comprising predominantly haemorrhage. Clusters B, D & F show enrichment for cell cycle and growth-related hallmarks, consistent with the ‘solid tumour’ histology evaluation. Cluster C shows enrichment of immune-related hallmarks and the Epithelial-Mesenchymal Transition hallmark, consistent with the presence of immune cell-marker proteins above. Cluster E shows enrichment for the “KRAS Signalling Dn” and “Epithelial-Mesenchymal Transition” hallmarks.

Clustering in high- and low-resolution maps was broadly consistent (Figure 5A) with generally contiguous clusters that represent the solid tumour (cluster 1), the brain/tumour interface (cluster 3), the margin between (clusters 2 & 5), and blood vessels (cluster 6). A volcano plot between cluster 3 and cluster 1 reflects the previously observed differential abundance of PYGL, ASPH and PRPH and other marker candidates (Figure 5B). Functional analysis of clusters 1 & 3 indicates that proteins involved in extracellular matrix, cell adhesion & motility, angiogenesis, immune processes, epidermis function, and neuronal development are differentially abundant between solid tumour and brain/tumour interface (Figure 5C).

**Figure 5.**
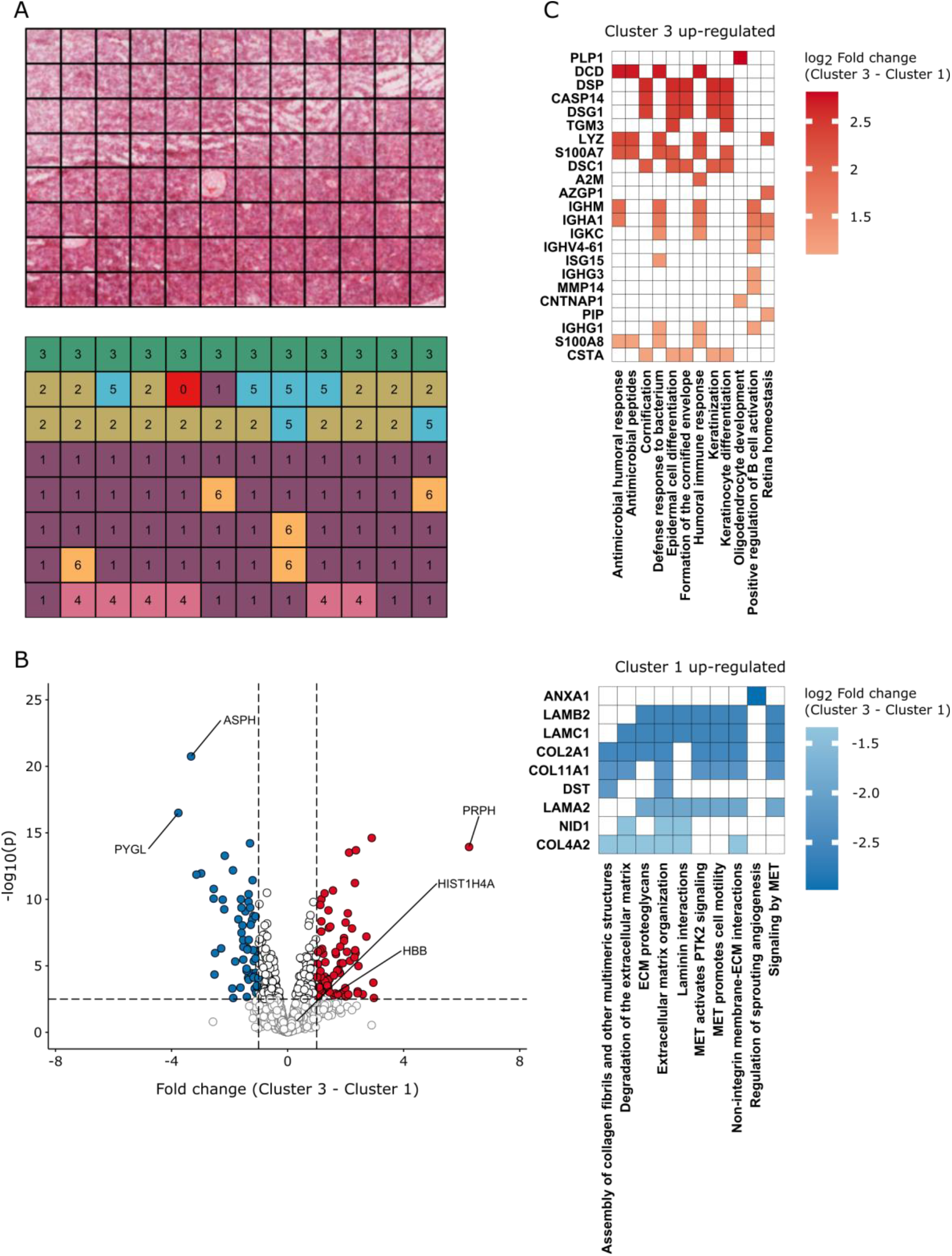
High-resolution proteomic map clustering defines functional layers at the tumour periphery. (A) Map of cluster assignment based on hierarchical clustering and the dynamic tree cut algorithm. Cluster 3 corresponds to brain/tumour interface. Cluster 1 corresponds to solid tumour. Clusters 2 & 5 correspond to the transition between solid tumour and brain/tumour interface (B) Volcano plot of clusters 3 against cluster 1 from (A). The horizontal dashed line represents an FDR of 5 %. The vertical dashed line represents -/+ 2-fold-change. (C) GSEA of clusters 1 & 3 from (A). Gene set membership is indicated by colouring the cell with that protein’s log_2_ fold-change.

We investigated the expression of other immune cell-marker proteins because of the above differences above in immune processes, and highly localised CD45 abundance and staining (Figure 3A). Two neutrophil markers, neutrophil cytosolic factor 2 & 4 (NCF2 & NCF4), show co-localisation with CD45 in the upper left of the sampled region along with two marker proteins for pro-tumour M2 macrophages, CD163 & mannose receptor C-type 1 (CD163 & MRC1)^40,41^. These proteins’ peak expression locations correspond with clusters 13 and 6 for the neutrophil and macrophage markers, respectively (Figure S4) and shows increased abundance for many proteins involved in neutrophil function and other immune-related processes such as B-cell differentiation and the JNK cascade within cluster 13. Within cluster 6, proteins involved in cell death, the cell cycle, morphogenesis, hedgehog signalling, collagen, and cytoskeletal organisation show increased abundance (Figure S5).

### MALDI imaging visualises lipid distributions consistent with proteomic maps

Due to the relatively coarse resolution of out proteomics approach, we also performed mass spectrometry imaging to investigate the spatial distribution of lipids within the tissue on an adjacent section using MALDI (Figure S6). Bisecting K-means clustering of the MALDI imaging pixels derived from 498 molecular ions broadly reflects the proteomic and cluster maps’ patterns, but with higher spatial resolution (20 µm). The upper-left region is a distinct cluster, and the boundary between the solid tumour and brain/tumour interface is visible. The ion image of haeme can be used to visualise the vascularisation of the tissue, and shows variability across the tissue, potentially indicating areas of nutrient gradients within the tissue.

Plotting the ion image of the 689.56 and 673.58 m/z ions, which correspond to the m/z of cholesterol ester^42^ (18:1) [M+K]^+^ and [M+Na]^+^ respectively, shows the highest intensity in the region of the tissue that corresponds to the area of high macrophage marker expression (cluster 6) in the proteomic maps (Figure S6), consistent with M1/M2 macrophage accumulation at the tumour interface visible in Figure S6 and Figures 2&3^43^.

### Spatial network clustering reveals extracellular matrix and integrin receptor heterogeneity

Throughout the spatial analysis and manual inspection of the data we noticed the common presence of many extracellular matrix-related proteins in the resulting enriched pathways and highly spatially-correlated proteins. Because of the relevance of the ECM within many cellular pathways and tumour development/progression, we performed ANF clustering based on the proteins annotated as ‘Core Matrisome’ proteins within MatrisomeDB^44^. This resulted in nine clusters (A-I) which are generally spatially contiguous across the tissue (Figure 6A). ECM-defined clusters were used for functional enrichment of proteins within each cluster, (Figure 6B) and are generally consistent with the functional enrichments in Figure 4E without ECM focus, linking overall spatial proteome distribution with ECM architecture in the tumour structure. To determine whether differential ECM abundance or composition was driving this clustering, we plotted the summed (Figure S7A) and mean (Figure S7B) abundance of the core matrisome proteins. These aggregate ECM spatial abundances show that both total ECM abundance (in the haemorrhage and immune infiltration area; clusters A & C in Figure 6A) and ECM composition (in the remaining clusters) are contributing to these cluster definitions.

**Figure 6.**
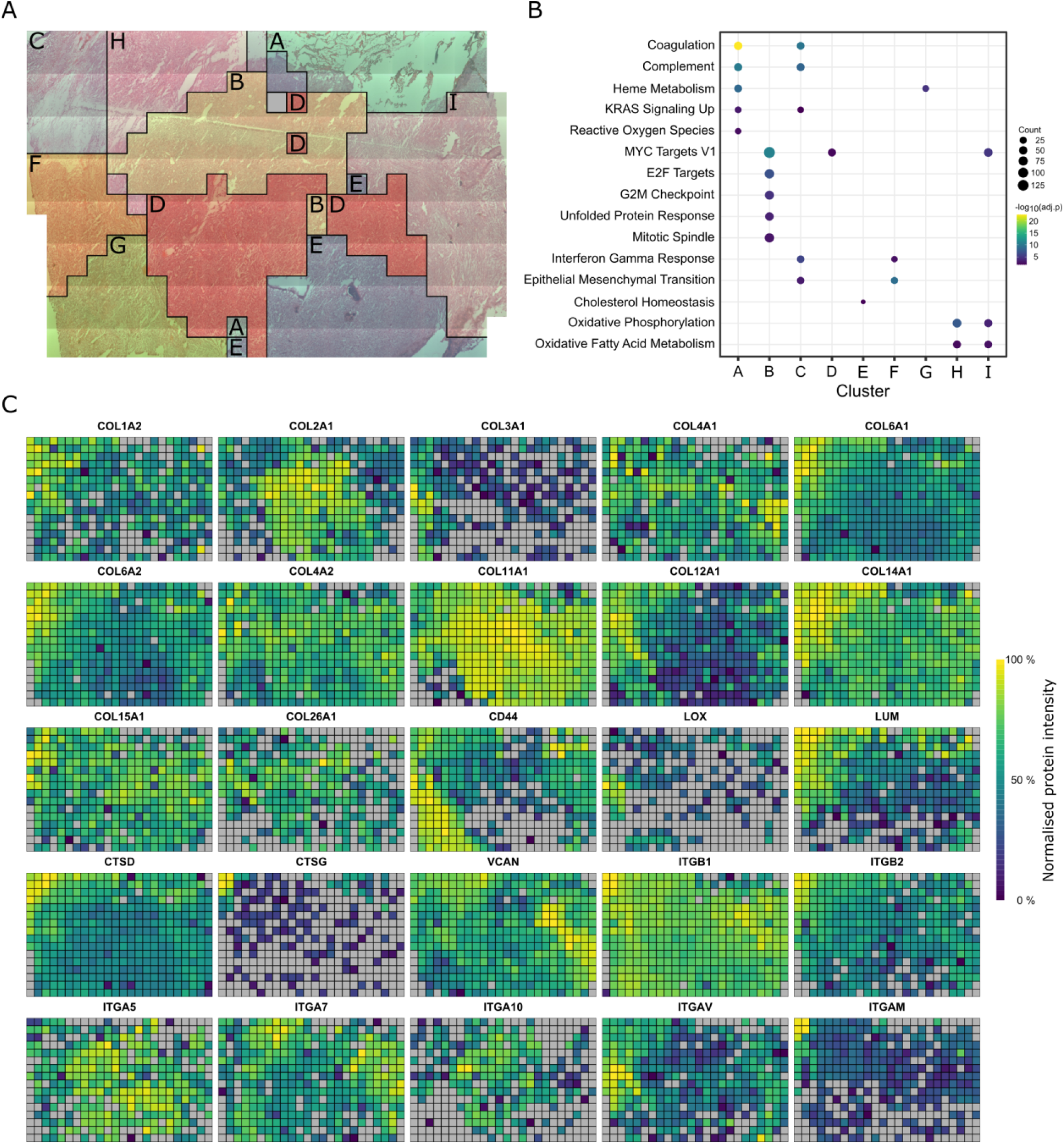
Spatial heterogeneity of extracellular matrix proteins across the ATRT tumour. (A) Cluster-map of ATRT tissue generated by affinity network fusion (ANF) of core matrisome proteins as defined in MatrisomeDB. This clustering is spatially-aware, meaning the relative spatial locations of samples and protein abundance values are used to generate these clusters. The grey pixel represents one sample where no core matrisome proteins were detected. (B) Enriched MSigDB Hallmark gene sets within marker proteins of the clusters shown in the ANF cluster-map. Significantly enriched hallmarks for each cluster are indicated by the presence of circles. The size and colour of the circles represent the number of proteins contributing to that term and the adjusted *p* value of the enrichment, respectively. (C) Proteomic maps of selected extracellular matrix proteins and integrin receptors at 833 µm resolution. Normalised protein intensities are scaled separately for each protein. Grey = not detected

Plotting the spatial distributions of detected collagens reveals broad heterogeneity in abundance of collagen proteins across the different collagen subfamilies (Figure 6C). The fibrillar collagens COL2A1, COL11A1, and COL11A2 show higher expression in the solid tumour region and other fibrillar collagens, COL1A2 and COL3A1, show higher abundance in the brain/tumour interface region. Other collagen subfamily members also show differential spatial abundance: COL6A1, COL6A2, COL26A1 (filament forming collagens); COL12A1, COL14A1 (fibril associated collagens). Collagens 4A1, 4A2 (network forming collagens), 15A1 and 18A1 (multiplexins) show spatially homogeneous expression. These spatial distributions broadly map to the ANF clusters. In addition, several integrin proteins also show spatial heterogeneity within the tissue (Figure 6C). Integrin α5, α7, α10, αV, αM and β2 show spatial heterogeneity whereas integrin β1 shows relatively homogeneous expression. Other proteins associated with the ECM also show spatial heterogeneity such as CD44, Lysyl oxidase (LOX), Lumican (LUM) (lower in solid tumour region), Versican (VCAN), and Cathepsin D & G (higher in immune cell-enriched region). In general ECM and ECM-associated proteins as defined in MatrisomeDB show a higher mean Moran’s I value than non-ECM proteins (ECM: I = 0.334, ECM-associated: I = 0.320, non-ECM: I = 0.219) (Figure S8)^44^.

## Discussion

The presented spatial proteomics workflow addresses the need to generate highly multiplexed, quantitative, spatially resolved measurements of proteins within tissue to understand the spatial organisation of molecular pathways in health and pathology. We demonstrate methods for the systematic sampling of tissue sub-sections using laser capture microdissection (LCM), sample preparation, liquid chromatography-mass spectrometry and advanced statistical analysis of topographic data. We have shown that these optimised methods are capable of detecting proteins and pathways with spatially variable abundance within tumour and tumour-infiltrated normal tissue. The application of spatially aware data analysis pinpoints processes with deep molecular resolution without prior knowledge of the tissue composition, thus creating an objective, unbiased way of deep phenotyping pathological tissue in its biological context.

A growing number of studies are currently focused on feature-driven LCM-coupled proteomics. In contrast, here, we propose a systematic approach, which allows the unbiased creation of comprehensive proteomic maps at the individual protein or pathway level.

These maps can fulfil the requirements for feature-driven analysis by reconstituting features from systematic sampling and allow the discovery of new proteo-phenotypes without the requirement of visually identifiable parameters.

Through our generation of proteomic maps, we have demonstrated the presence of molecular heterogeneity at multiple scales within ATRT tumour tissue sections, revealing proteomic differences between areas of tissue that appear visually homogeneous with good agreement with immunohistochemistry staining. Additionally, we demonstrate that this approach can also generate information on immune cell infiltration and state within the tissue, by the detection of neutrophil and pro-tumour M2 macrophage markers at different distances from the solid tumour.

We also have demonstrated that this approach provides spatial information on the cellular microenvironment across the tissue. The increased level of fibrillar collagens within the solid tumour is consistent with observations of increased fibrillar collagen deposition in glioblastoma tumours and the measurement of increased tissue stiffness of tumour types including brain tumours^45,46^. The spatial abundance of integrin subunits αM and β2 correlates with neutrophil marker abundance, strongly suggesting localised neutrophil accumulation^47^. Integrin α10 abundance correlates well with the abundance of COL2A1 and COL11A1, consistent with the observation that the α10β1 integrin receptor binds to type 2 and type 11 collagens^48^. Although functional interaction between these proteins cannot be determined from the methods used here, further investigation into integrin receptor signalling in ATRT could be a worthwhile avenue of investigation as integrin α10/β1 has shown potential as a therapeutic target in a glioblastoma xenograft model^49^.

The work we present here has several avenues for future improvement to generate more comprehensive information about biological structures. Integrating multiple data modalities and patient data is critical to a more comprehensive understanding of disease processes relying on advanced bioinformatics tools^50,51^. The same is expected to be true of spatially-resolved data. Further development of analytical and statistical methods that are aware of spatial relationships between samples is also required to maximise the utility of spatially resolved proteomics. Missing value imputation and machine learning approaches could benefit from taking samples’ spatial relationships into account^52,53^.

Spatial resolution should be balanced with the expectation for a meaningful depth to cover pathways of interest and technical limitations for sensitivity and throughput. Increasing the spatial resolution demonstrated here towards the single-cell level and covering comparable areas would be a formidable analytical challenge, further escalating when analysing tissue in three dimensions, despite the development of novel high-throughput LC-MS platforms that can now robustly analyse 1000s of samples relatively quickly^54–56^. Therefore, future approaches are likely to use systematic spatial proteomic analysis, possibly compromising spatial resolution but incorporating elements of machine learning, using orthogonal higher resolution ‘omics and imaging data to infer protein abundance towards individual cell resolution, as can be done on spatial transcriptomics data^57–60^. With the detection of LCM-based and cell-type resolved deep proteomes, these data will be highly complementary to current imaging technologies and increase the understanding of spatially resolved biological and pathological processes at the molecular level^24,31,61^.

## Acknowledgements

SD acknowledges support from the Nuffield Department of Medicine. SD, PDC, BMK & RF acknowledge support from the Chinese Academy of Medical Sciences Medical Sciences 2018-I2M-2-002. PDC was supported by Pfizer funding awarded to BMK.

We acknowledge the Oxford Brain Bank, supported by the Medical Research Council (MRC, MR/L022656/1) and Brains for Dementia Research (BDR) (Alzheimer Society and Alzheimer Research UK). This research project was funded by the NIHR Oxford Biomedical Research Centre (to OA, BRC-1215-20008). The views expressed are those of the authors and not necessarily those of the NHS, the NIHR, or the Department of Health. This work uses data provided by patients and collected by the NHS as part of their care and support and would not have been possible without access to this data. The NIHR recognises and values the role of patient data, securely accessed and stored, both in underpinning and leading to improvements in research and care.

## Methods

### Tissue retrieval and processing

Post-mortem brain tissue was retrieved by the Oxford Brain Bank; a research ethics committee (REC) approved and HTA regulated research tissue bank (REC reference 15/SC/0639). The retrieved brain was sectioned into 1 cm thick coronal sections starting at the level of the mammillary bodies. Due to its large size, tumour tissue was present within multiple of these coronal slices. The tumour tissue was dissected from the first coronal and second posterior coronal slices (P1 and P2). The tumour from the P2 slice was split into quadrants. Cryosections were taken from all pieces to determine the tissue block with the best morphological and cellular preservation. Cryosections were stained with H&E (see below) and examined by a Neuropathologist (OA). The upper-right quadrant from the P2 coronal slice was selected for use in further experiments.

Relevant tissue blocks of the AT/RT tumour were acclimatised to -20 °C and mounted onto a cryostat block using OCT Compound (Cell Path, ARG1180). Careful consideration was taken to ensure cut sections were not contaminated with OCT. Sections were cut at 10 µm and mounted onto UV irradiated (254 nm, 30 minutes) 1.0 PEN membrane slides (Zeiss) at -18 °C for LCM or Superfrost glass slides for histology. Sections were then air-dried for several minutes and placed onto a Shandon Linistain for automated H&E staining. Sections were fixed in 70 % denatured alcohol, hydrated, stained with Harris’ Haematoxylin, incubated in 0.4 % acid alcohol, placed in Scot’s tap water, and stained with Eosin containing 0.25 % acetic acid with regular washing steps in between. Stained sections were then dehydrated in increasing concentrations of denatured alcohol and air-dried without coverslips and stored at -80 °C until processing by laser-capture microdissection.

### Laser capture microdissection

Areas of tissue analysed were annotated and isolated from the prepared slides using a laser-capture microscope equipped with laser pressure catapulting (PALM Microbeam, Zeiss). Cutting and capturing the annotated tissue areas were performed automatically and used the 10x objective lens. The settings in the control software for cutting were Energy: 43, Focus: 55; and for capturing were Energy 20, Focus -15. Samples were collected into 20 µL RIPA buffer (Pierce #89900) in the cap of 200 µL PCR tubes or PCR-cap strips of 8. Collected samples were immediately placed in dry ice. Samples were stored at -80 °C until further use.

### Proteomic sample processing

Samples were thawed, incubated at room temperature for 30 minutes and briefly centrifuged. Caps were rinsed with 20 µL of RIPA buffer (#89900, Pierce) containing 25 units of Benzonase (E1014, Merck) to collect any remaining tissue and briefly centrifuged, followed by incubation at room temperature for 30 minutes to degrade DNA and RNA. Proteins were reduced by adding DTT to 5 mM and incubated at room temperature for 30 minutes, followed by the addition of iodoacetamide to 20 mM and incubation at room temperature for 30 minutes.

Paramagnetic SP3 beads (GE45152105050250 & GE65152105050250, Cytiva) were prepared as described by Hughes *et al*. and processed by a modified SP3 protocol ^24,62,63^. Three µL of SP3 beads were mixed with the samples, and acetonitrile added to a final concentration of 70 % (v/v). Samples were mixed with 1000 rpm orbital shaking for 18 minutes, followed by bead immobilisation on a magnet for 2 minutes. The supernatant was discarded, and beads were washed twice with 70 % (v/v) ethanol in water and once with 100 % acetonitrile without removal from the magnet. Beads were resuspended in 50 mM ammonium bicarbonate containing 25 ng of Trypsin (V5111, Promega) and digested overnight at 37 °C. After digestion, the beads were resuspended by bath sonication. Acetonitrile was added to the samples to 95 % (v/v) and shaken at 1000 rpm for 18 minutes. Beads were immobilised on a magnet for 2 minutes, and the supernatant discarded. Beads were resuspended in 2 % acetonitrile and immobilised on a magnet for 5 minutes. Peptides were transferred to glass LC-MS vials or 96-well PCR plates containing formic acid in water, resulting in a final formic acid concentration of 0.1 %.

### LC-MS/MS

Peptides from 833 µm resolution samples were analysed by LC-MS/MS using a Dionex Ultimate 3000 (Thermo Scientific) coupled to a timsTOF Pro (Bruker) using a 75 μm x 150 mm C18 column with 1.6 μm particles (IonOpticks) at a flow rate of 400 nL/min. A 17-minute linear gradient from 2 % buffer B to 30 % buffer B (A: 0.1 % formic acid in water. B: 0.1 % formic acid in acetonitrile) was used^64^. The TimsTOF Pro was operated in PASEF mode.

The TIMS accumulation and ramp times were set to 100 ms, and mass spectra were recorded from 100 – 1700 m/z, with a 0.85 – 1.30 Vs/cm^2^ ion mobility range. Precursors were selected for fragmentation from an area of the full TIMS-MS scan that excludes most ions with a charge state of 1+. Those selected precursors were isolated with an ion mobility dependent collision energy, increasing linearly from 27 – 45 eV over the ion mobility range. Three PASEF MS/MS scans were collected per full TIMS-MS scan, giving a duty cycle of 0.53 s. Ions were included in the PASEF MS/MS scan if they met the target intensity threshold of 2000 and were sampled multiple times until a summed target intensity of 10000 was reached. A dynamic exclusion window of 0.015 m/z by 0.015 Vs/cm^2^ was used, and sampled ions were excluded from reanalysis for 24 seconds.

Peptides from 350 µm resolution samples were analysed by nano-UPLC-MS/MS using a Dionex Ultimate 3000 coupled to an Orbitrap Fusion Lumos (Thermo Scientific) using a 75 µm x 500 mm C18 EASY-Spray Columns with 2 µm particles (Thermo Scientific) at a flow rate of 250 nL/min. A 60-minute linear gradient from 2 % buffer B to 35 % buffer B (A: 5 % DMSO, 0.1 % formic acid in water. B: 5 % DMSO, 0.1 % formic acid in acetonitrile). MS1 scans were acquired in the Orbitrap between 400 and 1500 m/z with a resolution of 120,000 and an AGC target of 4 × 10^5^. Precursor ions between charge state 2+ and 7+ and above the intensity threshold of 5 × 10^3^ were selected for HCD fragmentation at a normalised collision energy of 28 %, an AGC target of 4 × 10^3^, a maximum injection time of 80 ms and a dynamic exclusion window of 30 s. MS/MS spectra were acquired in the ion trap using the rapid scan mode.

### Proteomic data analysis

Raw data files were searched against the UniProtKB human database (Retrieved 17/01/2017, 92527 sequences) using MaxQuant version 1.6.14.0, allowing for tryptic specificity with up to 2 missed cleavages. Cysteine carbamidomethylation was set as a fixed modification. Methionine oxidation and protein N-terminal acetylation were set as variable modifications and the “match between runs (MBR)” option was used (MBR was not used for tissue titration data). All other settings were left as default. Label-free quantification was performed using the MaxLFQ algorithm within MaxQuant^65,66^. Protein and peptide false discovery rate (FDR) levels were set to 1 %.

### Spatial data analysis

The mapping of the mass spectrometry raw files to their relative spatial locations is shown in Supplementary Table 1. The spatial analysis uses functions within the spdep and raster R packages^67–69^. MaxQuant’s protein level output files (‘proteingroups.txt’) were filtered to remove reverse hits, ‘Only identified by site’ hits and potential contaminants. The ‘LFQ intensity’ columns were log_2_ transformed and then normalised by median subtraction.

Protein groups that did not meet a cut-off of having at least 9 pixels with normalised LFQ values are not taken forward for further analysis. The following steps occur independently for each protein group. Normalised LFQ intensities were then coerced into a matrix reflecting the rastered pattern of sample acquisition. The quantification matrix was converted into a raster object and then to a polygon object using the raster R package. From this polygon object, a neighbour list was built for each pixel of the raster using the ‘Queen’s Case’ where cells are considered neighbours if they share an edge or a vertex. The neighbour list was then supplemented with a spatial weights matrix using a binary coding scheme where neighbours are given a weighting of ‘1’ and non-neighbours a weighting of ‘0’ in the spatial weights matrix. The raster object and the weighted neighbour list were then used as inputs to a permutation test for the Moran’s *I* statistic, calculated using 999 random spatial permutations of the raster object to calculate pseudo-*p*-values. Moran’s *I* statistics and the associated *p*-values are collected for every protein group. The *p*-values were then corrected for multiple testing using the Benjamini-Hochberg FDR method.

### Immunohistochemistry

Sections were cut as above and were mounted to Superfrost glass slides for IHC and air-dried. Slides were fixed in ice-cold acetone for 10 minutes, washed twice with TBS/T (20 mM Tris, 150 mM NaCl, 0.05 % Tween 20) and blocked with 10 % goat serum in TBS/T for 60 minutes at room temperature. Primary antibodies were diluted in 5 % goat serum in TBS/T and incubated at RT for 60 minutes or 4 °C overnight. Sections were washed three times with TBS/T. Staining visualisation was performed by incubating with a cocktail of anti-mouse and anti-rabbit secondary antibodies conjugated to horseradish peroxidase (Envision Kit, Agilent) for 60 minutes at room temperature. Sections were then washed with TBS/T three times and incubated with 2 % 3,3’-diaminobenzidine for 5 minutes, immersed in water and then counterstained with Harris’ Haematoxylin for 1 minute. Primary antibodies used and dilutions: rabbit anti-PYGL, 1:100, 4 °C overnight, HPA000962 (Atlas Antibodies); rabbit anti-ASPH, 1:1000, RT 60 minutes, NBP2-34125 (Novus Biologicals); mouse anti-CD45 (PD7/26 + 2B11), 1:200, RT 60 minutes, ab781 (Abcam).

### Clustering

Empty pixels and pixels covering the large region of haemorrhage were not included. A distance matrix was built containing the Euclidean distance between each pixel’s set of protein LFQ values. Hierarchical clustering of the distance matrix was performed in R using the “average” agglomeration method. Dendrograms were cut using the Dynamic Tree Cut method at a height setting of 100^70^.

A complete input matrix is required for UMAP visualisation^71^, so proteins with fewer than 70 % valid values across the experiment were removed. The remaining missing values were imputed in on a per-sample basis by random draws from a normal distribution using a width of 0.3 and a downshift of 1.8. UMAP dimensionality reduction was performed on this imputed data with default settings, and the first two embedding components plotted, and samples coloured according to their cluster assigned by Dynamic Tree Cut at a height of 100.

### Pathway Analysis

A one-way ANOVA test was performed in R to test for differences in means between the clusters generated by Dynamic Tree Cut at a height of 100. A pairwise post-hoc correction was applied using Tukey’s Honestly Significant Difference method. The resulting pairwise comparisons were used as inputs to ClusterProfiler’s gene set enrichment analysis to test for enrichment of Gene Ontology Biological Process terms using a Fisher’s exact test with a 5 % FDR threshold^72^.

### Lipid MALDI imaging

Vacuum dried sections were scanned with a TissueScout scanner at 3200 dpi (Bruker) to generate a reference image for later position teaching. Dihydroxybenzoic acid (DHB) was dissolved at a concentration of 15 mg/ml in 90 % ACN, 0.1 % TFA and sprayed on top of the AT/RT sections using a TM-sprayer (HTX Technologies). The matrix was applied in a criss-cross pattern with 3 mm track spacing at a 1200 mm/min nozzle velocity. Fourteen layers were sprayed at a 0.125 ml/ml flow rate using 50 % ACN as the liquid phase at 10 psi pressure. The nozzle temperature was set to 60 °C and the distance of the nozzle to the section was 40 mm. All imaging data were acquired on a timsTOF fleX instrument (Bruker) which is equipped with a dual ESI and MALDI source in positive Q-TOF mode. External calibration was performed using red phosphorous which was spotted next to the section. The laser was operated in beam scan mode, ablating an area of 15×15 µm resulting in a pixel size of 20 µm. The repetition rate of the laser was set to 10 kHz and 400 laser shots were acquired per pixel. Data were acquired in the mass range 300-1400 m/z.

The software SCiLS Lab (version 2020a; Bruker) was used for MALDI Imaging data analysis. All data were root-mean-square (RMS) normalised. After importing the data, an unsupervised segmentation was calculated using the bisecting k-means algorithm and a peak list containing 302 m/z-intervals with correlation used as a distance metric^73^. The resulting segmentation map was split into several clusters that resemble the histopathology of the tumour section. The identification of cholesterol ester, observed as 689.5607 and 673.5843 m/z ions, which correspond to the m/z adducts of cholesterol ester (18:1) [M+K]^+^ and [M+Na]^+^, was performed using LIPID-MAPS database searches using exact mass^74^.

Expected m/z were 689.5633 (M+K^+^), 673.5894 (M+Na^+^) and delta masses were 0.0026 (3.7ppm) and 0.0051 m/z (7.5ppm), respectively. The next closest match corresponding to a species with both Na^+^ and K^+^ adducts is DG 38:1 with delta masses of 0.0102 (M+Na^+^) and 0.0126 (M+K^+^) m/z.

### Affinity Network Fusion

For the network fusion approach, we used a minimally processed expression matrix, log2 transformed and median centred, removing all empty pixels. We selected the top 25% of proteins by variance across all pixels and used this to calculate a proteomic similarity matrix of Spearman’s rank correlation coefficients. For the focussed extracellular matrix protein analysis, all detected proteins annotated as “Core Matrisome” in MatrisomeDB were used to calculate the protein similarity matrix. We converted this to a proteomic distance matrix (by taking 1-similarity). Separately, we created a complementary spatial distance matrix representing the Euclidean distance from each pixel location to each other pixel location (where horizontally and vertically adjacent neighbours are distance 1, diagonal neighbours are distance √2 and so on). We converted both matrices into affinity matrices and fused them using Affinity Network Fusion^38^. We then performed spectral clustering on the fused affinity matrix, where the number of clusters (6) was selected using the maximal eigengap heuristic.

## Data availability

The mass spectrometry proteomics data have been deposited to ProteomeXchange Consortium via the PRIDE partner repository and will be made available to reviewers upon submission.

## Supplementary Figures

**Figure S1.**
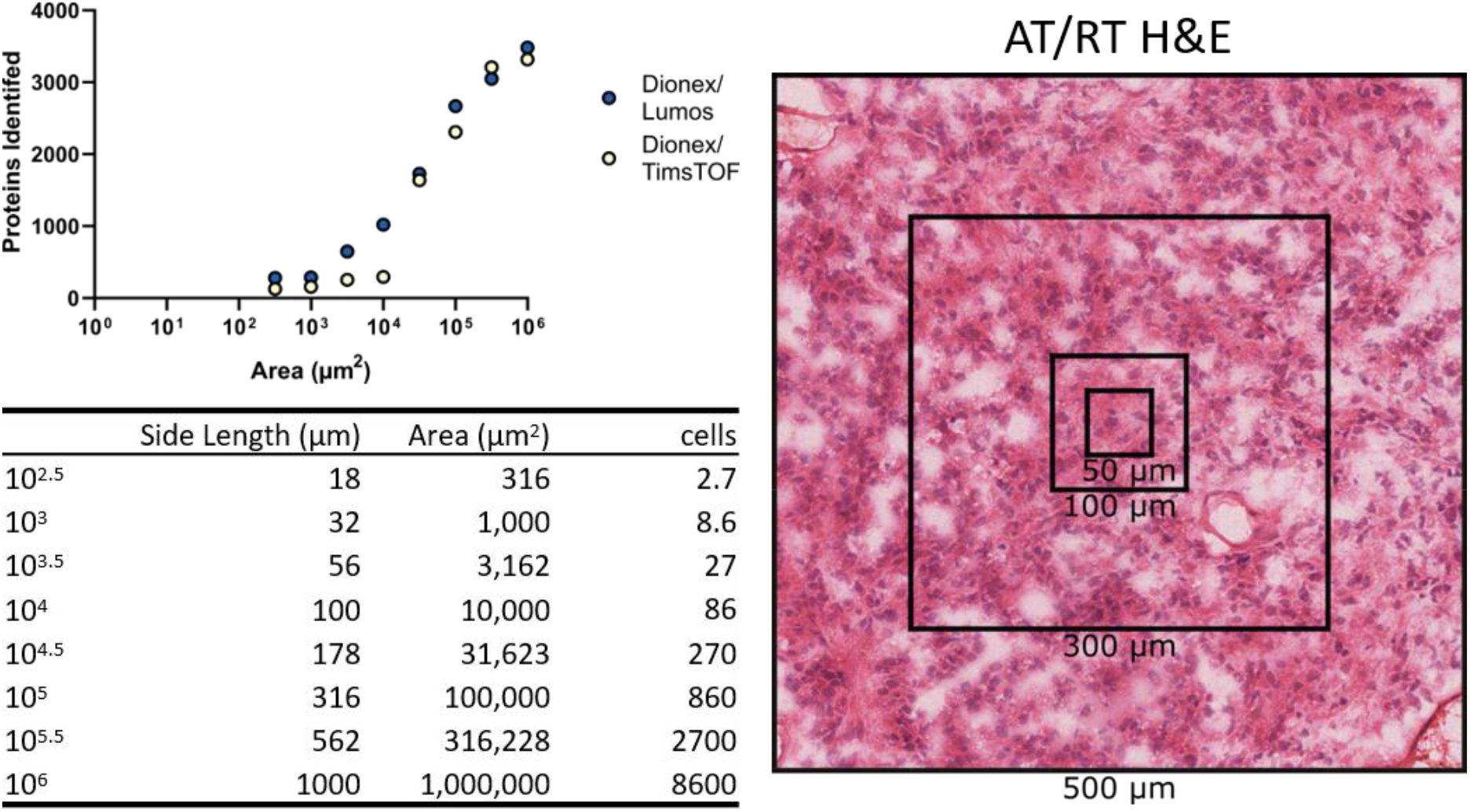
Tissue area titration across two LC-MS/MS systems. The number of proteins identified from a titration of AT/RT tissue area on two LC-MS/MS systems.

**Figure S2.**
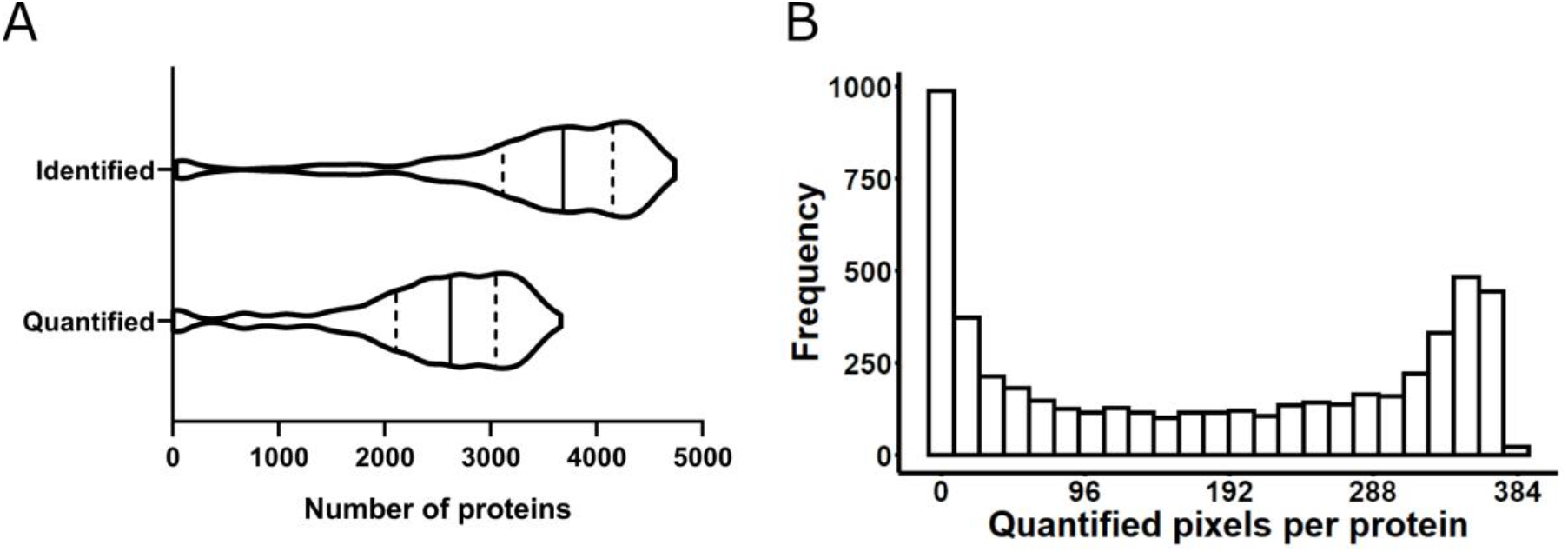
Distribution of identified and quantified proteins at 833 µm resolution. (A) Violin plots showing distributions of identified and quantified proteins per pixel. Solid vertical lines represent the median value. Dashed vertical lines represent upper and lower quartiles. (B) Histogram of quantified pixels per protein map.

**Figure S3.**
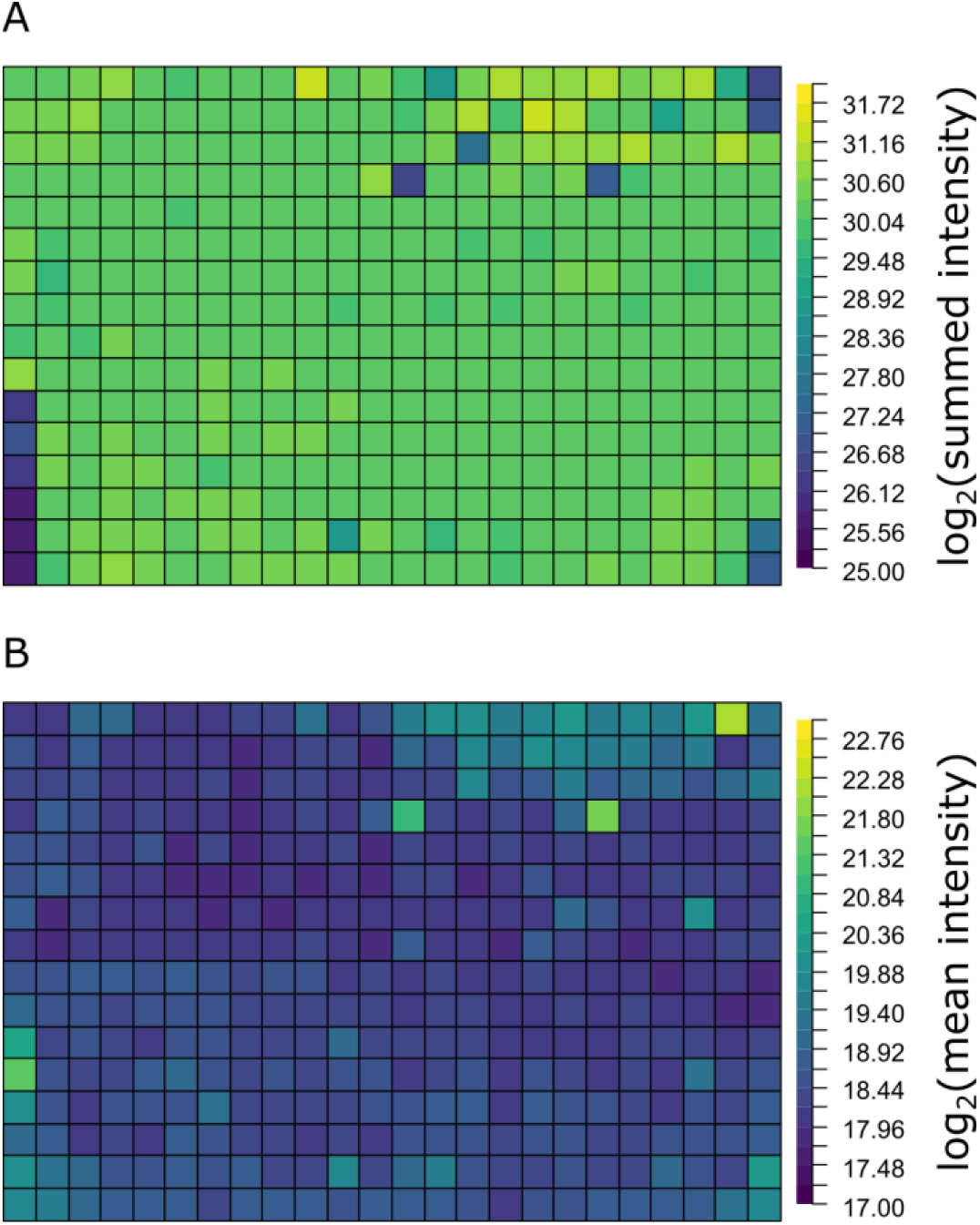
Aggregate intensity distributions. Maps of the log_2_ transformed (A) summed and (B) mean intensities of each pixel.

**Figure S4.**
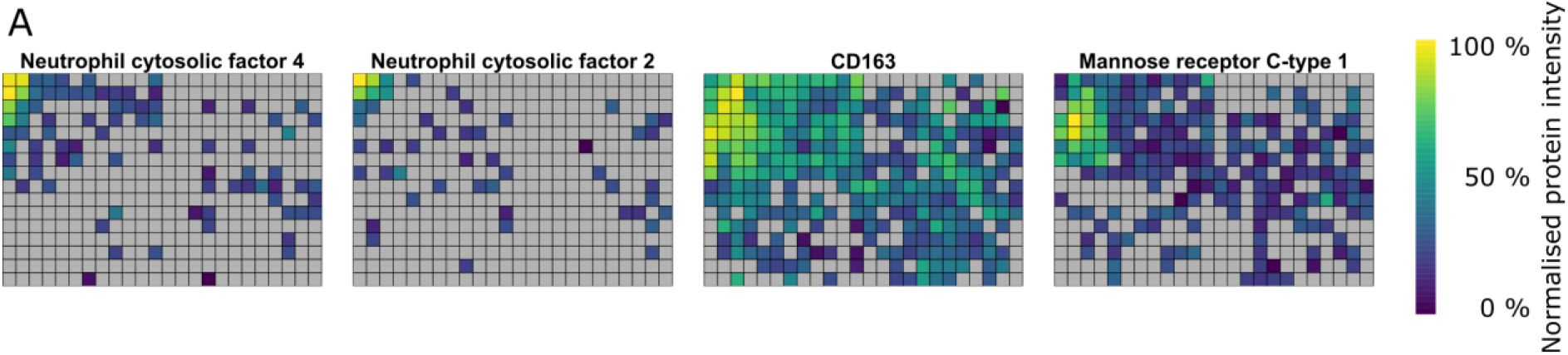
Proteomic maps of immune cell markers. Proteomic maps of immune cell-marker proteins at 833 µm resolution. Normalised protein intensities are scaled separately for each protein. Grey = not detected.

**Figure S5.**
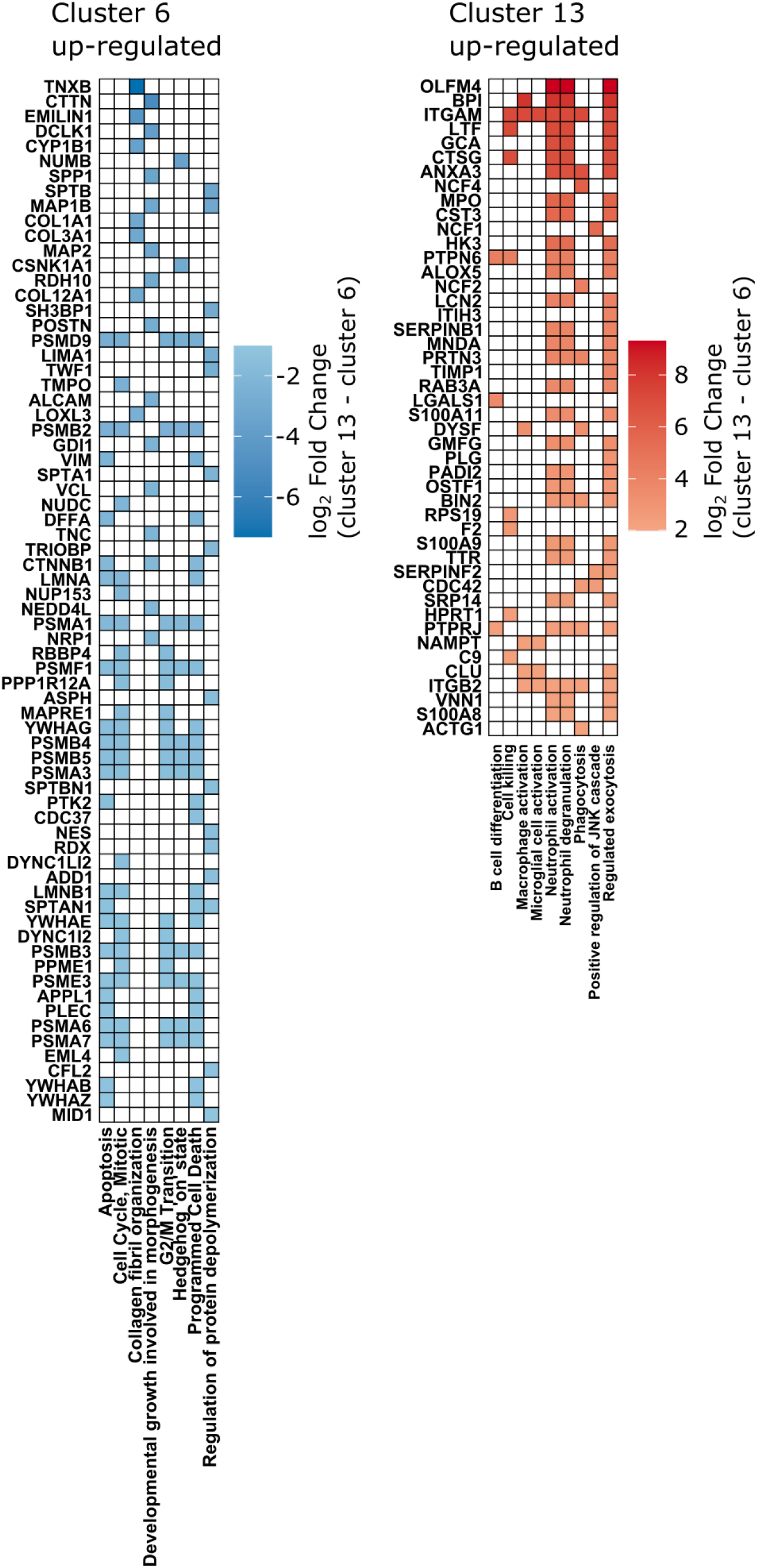
Pathway enrichment in immune-cell clusters. GSEA of clusters 6 & 13 from Figure 4A. Gene set membership is indicated by colouring the cell with that protein’s log2 fold-change.

**Figure S6.**
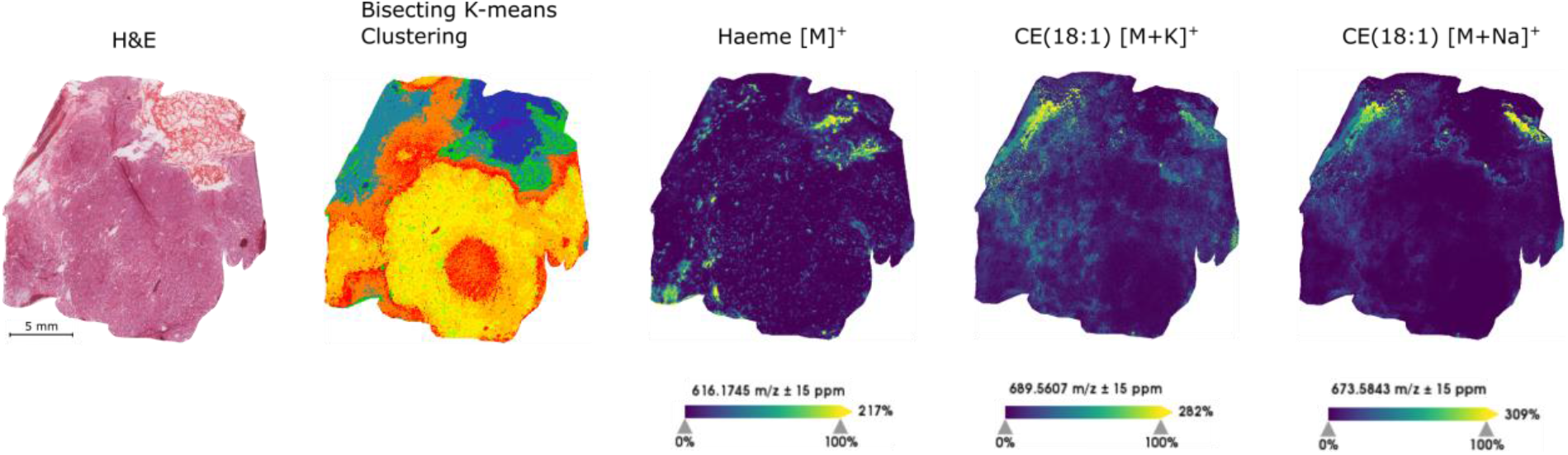
AT/RT brain tissue MALDI imaging shows regional enrichment of haeme and lipids at 20 µm resolution. (Left) Image of haematoxylin and eosin-stained tissue after MALDI acquisition. (Middle-left) MALDI Imaging pixels were clusters by Bisecting K-Means clustering and the generated clusters are indicated by colour. m/z images for ions corresponding to the masses of haeme (middle), cholesterol ester (18:1) [M+K]^+^ (middle-right) and [M+Na]^+^ (right) ions. Ion intensities are scaled separately for each ion.

**Figure S7.**
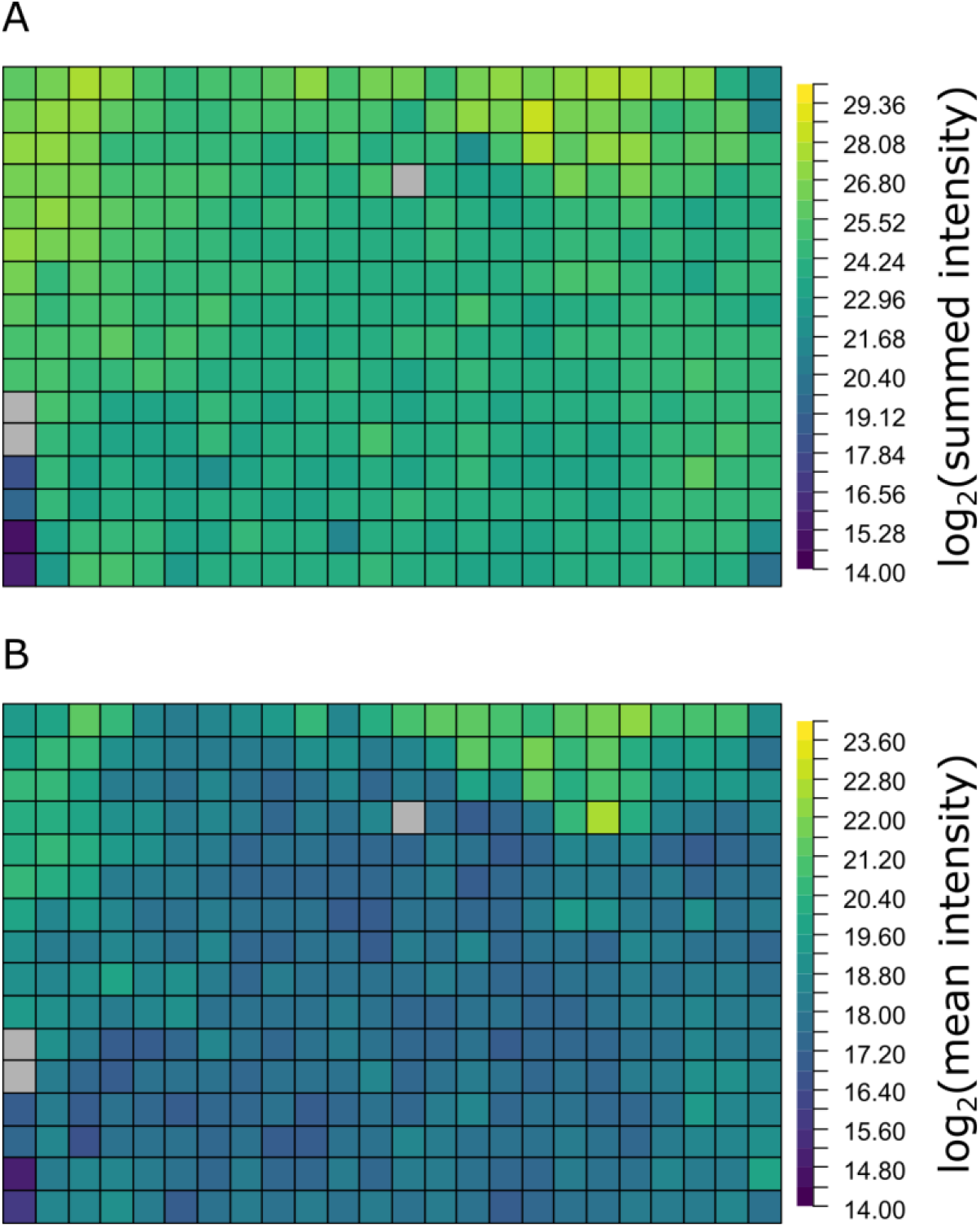
Aggregate intensity distribution of core matrisome proteins. Maps of the log_2_ transformed (A) summed and (B) mean intensities of core matrisome proteins as defined by MatrisomeDB for each pixel. Grey pixels indicate no core matrisome proteins were detected.

**Figure S8.**
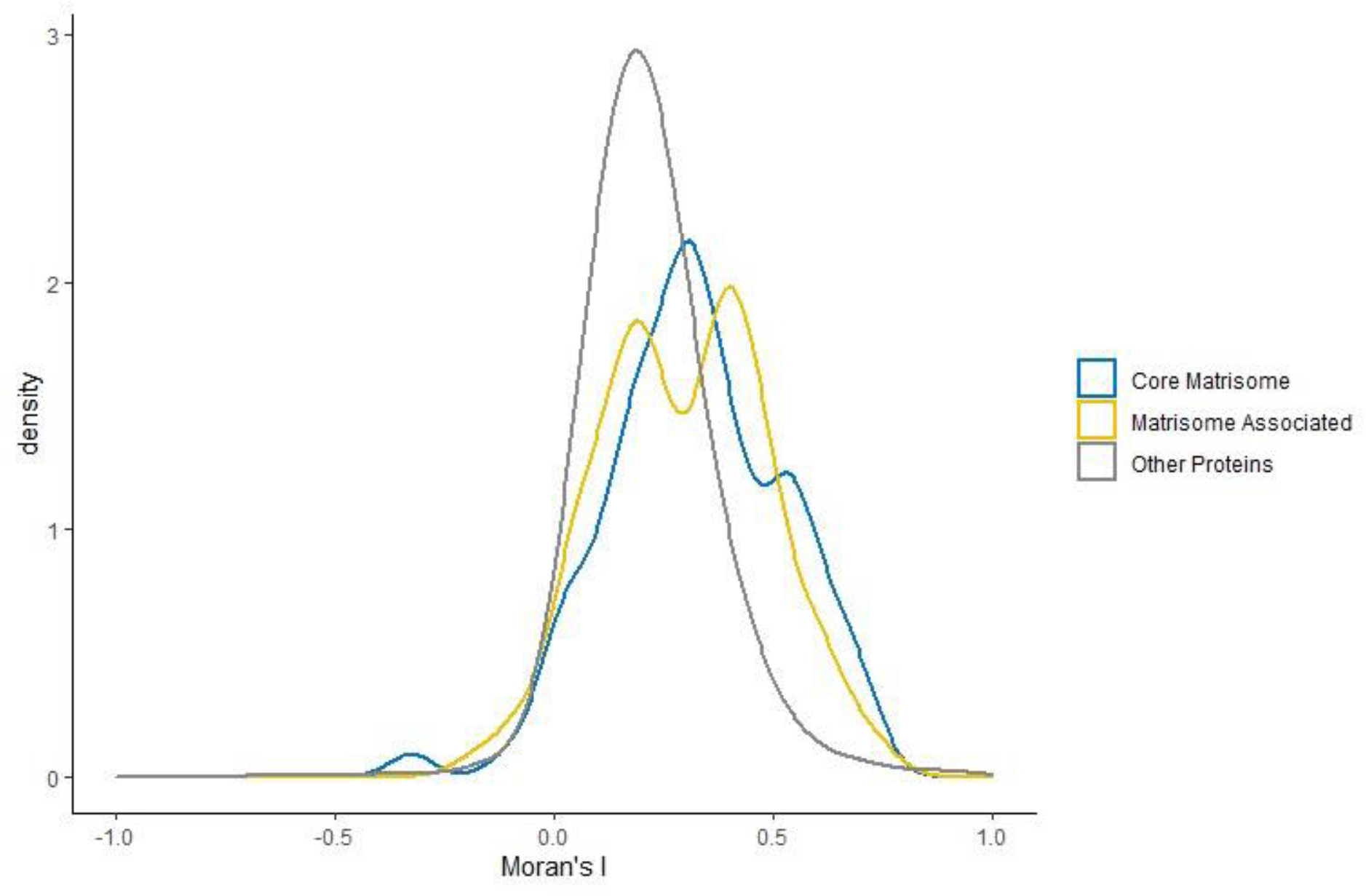
Distribution of Moran’s I in matrisome and non-matrisome proteins. Moran’s *I* distribution of proteins annotated in Matrisome DB as core matrisome (blue), matrisome associated (yellow), and all other proteins (grey).

